# Prdm8 regulates pMN progenitor specification for motor neuron and oligodendrocyte fates by modulating Shh signaling response

**DOI:** 10.1101/2020.03.27.012294

**Authors:** Kayt Scott, Rebecca O’Rourke, Austin Gillen, Bruce Appel

## Abstract

Spinal cord pMN progenitors sequentially produce motor neurons and oligodendrocyte precursor cells (OPCs). Some OPCs differentiate rapidly as myelinating oligodendrocytes whereas others remain into adulthood. How pMN progenitors switch from producing motor neurons to OPCs with distinct fates is poorly understood. pMN progenitors express *prdm8*, which encodes a transcriptional repressor, during motor neuron and OPC formation. To determine if *prdm8* controls pMN cell fate specification, we used zebrafish as a model system to investigate *prdm8* function. Our analysis revealed that *prdm8* mutant embryos have a deficit of motor neurons resulting from a premature switch from motor neuron to OPC production. Additionally, *prdm8* mutant larvae have excess oligodendrocytes and a concomitant deficit of OPCs. Notably, pMN cells of mutant embryos have elevated Shh signaling coincident with the motor neuron to OPC switch. Inhibition of Shh signaling restored the number of motor neurons to normal but did not rescue the proportion of oligodendrocytes. These data suggest that Prdm8 regulates the motor neuron-OPC switch by controlling the level of Shh activity in pMN progenitors and also regulates allocation of oligodendrocyte lineage cell fates.

**Summary Statement:** Prdm8 regulates the timing of a motor neuron-oligodendrocyte switch and oligodendrocyte lineage cell identity in the zebrafish spinal cord.

## INTRODUCTION

Oligodendrocytes, one of the major glial cell types of the central nervous system (CNS) of vertebrate animals, increase the speed of axon electrical impulses and support neuron health by wrapping axons with myelin membrane (Simons and Nave, 2016). In the spinal cord most oligodendrocytes originate from ventral pMN progenitors (Noll and Miller, 1993; Warf et al., 1991), which express the basic helix loop helix (bHLH) transcription factor Olig2 (Lu et al., 2000; Novitch et al., 2001; Zhou and Anderson, 2002; Zhou et al., 2000). pMN progenitors first produce motor neurons followed by oligodendrocyte precursor cells (OPCs) (Richardson et al., 2000; Rowitch, 2004). After specification, some OPCs rapidly differentiate as myelinating oligodendrocytes whereas other OPCs persist into adulthood (Dawson et al., 2003; Pringle et al., 1992). The switch from motor neuron to OPC production and subsequent regulation of oligodendrocyte differentiation require tight control of gene expression through a complex network of interacting transcription factors and extracellular cues. Whereas many factors that promote oligodendrocyte differentiation and myelination have been identified (Elbaz and Popko, 2019; Emery, 2010; He and Lu, 2013; Sock and Wegner, 2019; Zuchero and Barres, 2013), the mechanisms that regulate the onset of OPC specification and maintain some in a non-myelinating state are not well understood.

During early neural tube patterning, pMN progenitors are specified by the morphogen Sonic Hedgehog (Shh). The Shh ligand, secreted by notochord, a mesodermal structure below the ventral spinal cord, and floor plate, the ventral-most cells of the neural tube, signals neural progenitors to acquire ventral identities (Echelard et al., 1993; Martí et al., 1995; Roelink et al., 1994). The Shh ligand binds to its transmembrane receptor Patched (Ptch), which allows intercellular Shh signaling to be transduced by Smoothened (Smo). Upon Shh binding, Smo is internalized to promote GliA transcriptional activity by inhibiting its cleavage to GliR (Briscoe and Thérond, 2013; Danesin and Soula, 2017; Ribes and Briscoe, 2009). Graded Shh activity induces expression of genes that encode bHLH and homeodomain proteins at distinct positions on the dorsoventral axis (Briscoe and Thérond, 2013; Briscoe et al., 2000; Poh et al., 2002). The duration of Shh signaling also influences cell fate and gene expression. Initially, high ventral Shh signaling activates expression of Olig2, then sustained Shh activity promotes expression of Nkx2.2 adjacent to the floor plate, ventral to Olig2, thus forming two distinct ventral progenitor domains (Dessaud et al., 2007; Dessaud et al., 2010). The sequential induction of cross-repressive transcription factors by graded morphogen activity along the dorsoventral axis creates spatially restricted progenitor domains that sequentially give rise to specific neurons and glia (Briscoe et al., 2000; Lek et al., 2010; Nishi et al., 2015).

Shh activity is necessary for the establishment of the pMN domain and motor neuron formation and subsequently, a transient increase in Shh activity coincides with and is required for timely OPC specification (Danesin and Soula, 2017). Pharmacological and genetic reduction of Shh signaling in chick, mouse and zebrafish spinal cords resulted in prolonged motor neuron formation and impaired OPC formation (Al Oustah et al., 2014; Danesin et al., 2006; Hashimoto et al., 2017; Jiang et al., 2017; Ravanelli and Appel, 2015; Touahri et al., 2012). Further, chick neural tube explants treated with exogenous Shh led to premature formation of OPCs at the expense of motor neurons (Danesin et al., 2006; Orentas et al., 1999). Thus, transient elevation of Shh activity is required to induce the transition from motor neuron to OPC production. The temporal change in Shh activity is in part due to upregulation of Sulfatase 1/2 by p3 cells, which are ventral to pMN progenitors, prior to OPC specification (Al Oustah et al., 2014; Danesin et al., 2006; Jiang et al., 2017). Sulfatase expression increases local Shh ligand concentration available to pMN progenitors (Al Oustah et al., 2014; Danesin et al., 2006) and loss of Sulfatase 1/2 functions delays the motor neuron-OPC switch (Jiang et al., 2017). Whether additional mechanisms contribute to modulation of Shh signaling strength to regulate fate specification over time is not well understood.

In addition to expressing distinct combinations of bHLH and homeodomain transcription factors, subsets of spinal cord progenitors express specific PRDI-BF1 and RIZ homology domain containing (Prdm) proteins (Zannino and Sagerström, 2015). This family of proteins contain a N-terminal SET domain followed by a varied number of C-terminal zinc finger repeats and act as transcriptional regulators or co-factors implicated in nervous system patterning, neural stem cell maintenance and differentiation (Baizabal et al., 2018; Chittka et al., 2012; Hanotel et al., 2014; Hernandez-Lagunas et al., 2011; Kinameri et al., 2008; Ross et al., 2012; Thélie et al., 2015; Yildiz et al., 2019). In the ventral mouse spinal cord, neural progenitors express Prdm8 from E9.5 to E13.5 (Kinameri et al., 2008; Komai et al., 2009), corresponding to the period of motor neuron and OPC formation. The function of Prdm8 in spinal cord development is not yet known, but in the mouse telencephalon Prdm8 forms a repressive complex with Bhlhb5, a bHLH transcription factor closely related to Olig2, to regulate axonal migration (Ross et al., 2012). Moreover, in the retina Prdm8 promotes formation of a subset of rod bipolar cells and regulates amacrine cell type identity (Jung et al., 2015). These data raise the possibly that Prdm8 regulates pMN cell development.

To investigate *prdm8* expression and function in pMN progenitors we used the developing zebrafish spinal cord as a model. Our expression analysis showed that pMN progenitors express *prdm8* prior to and during the switch from motor neuron to OPC production and that, subsequently, differentiating oligodendrocytes downregulate *prdm8* expression. Because Prdm8 can control cell fate, we therefore hypothesized that Prdm8 regulates motor neuron and OPC specification. To test this hypothesis, we performed a series of experiments to assess changes in pMN cell fates resulting from loss of *prdm8* function. Our data reveal that *prdm8* mutant embryos have a deficit of late-born motor neurons, excess differentiating oligodendrocytes and a deficit of OPCs. Birthdating studies showed that the motor neuron deficit results from a premature switch from motor neuron to OPC production. *prdm8* mutant embryos have abnormally high levels of Shh signaling and pharmacological suppression of Shh signaling rescued the motor neuron deficit but not the formation of excess oligodendrocytes. Taken together, our data suggest that Prdm8 dampens Shh signaling activity to modulate the timing of the motor neuron-OPC switch and secondarily regulates the myelinating fate of oligodendrocyte lineage cells.

## RESULTS

### pMN progenitors and oligodendrocyte lineage cells express *prdm8*

To begin our investigation of *prdm8* function, we first assessed the temporal and spatial features of *prdm8* expression in the zebrafish spinal cord during development. To do so, we performed in situ RNA hybridization (RNA ISH) using transverse sections obtained from *Tg(olig2:EGFP*) embryos and larvae. pMN cells in these fish express EGFP driven by *olig2* regulatory DNA (Shin et al., 2003). pMN cells and cells dorsal to the pMN domain expressed *prdm8* at 24, 36 and 48 hours post-fertilization (hpf) (Fig. 1A). This is consistent with previous data that revealed that cells of the pMN, p2 and p1 domains in the developing mouse spinal cord express *Prdm8* (Kinameri et al., 2008; Komai et al., 2009). Next we evaluated pMN cell expression of *prdm8* through development within a single cell RNA-seq (scRNA-seq) data set obtained from *olig2*:EGFP^+^ cells isolated from 24, 36 and 48 hpf embryos, a period spanning formation of most motor neurons and OPCs. An aligned Harmony clustering analysis and Uniform Manifold Approximation and Projection (UMAP) of the data revealed that gene expression profiles formed several distinct clusters (Fig 1B). Plotting individual gene expression profiles revealed that many *olig2*^+^ *sox19a*^+^ cells, which likely represent pMN progenitors, also expressed *prdm8* (Fig. 1C-E).

**Fig. 1.**
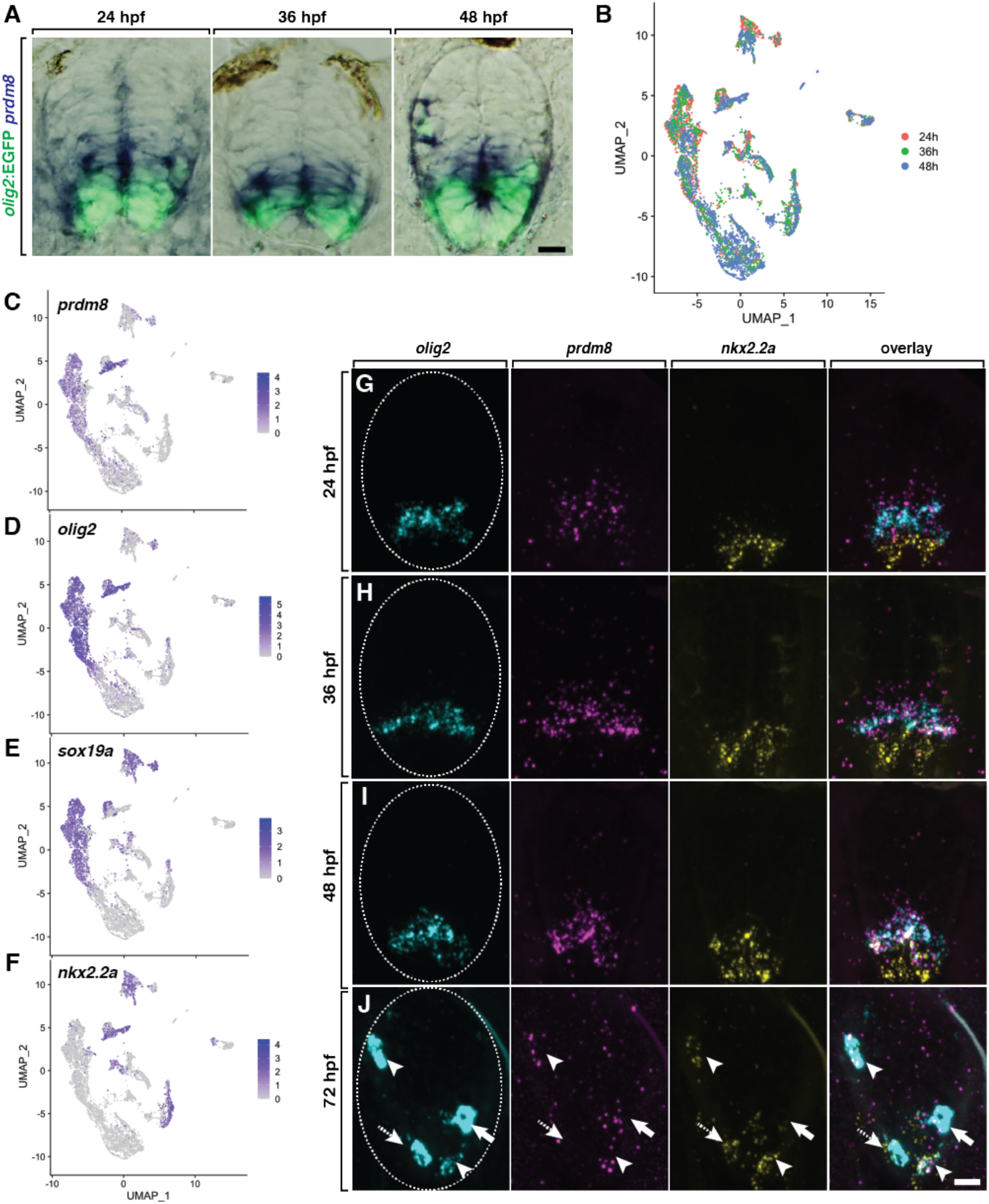
pMN progenitors express *prdm8*. (A) Representative transverse sections of trunk spinal cord, with dorsal up, showing *prdm8* RNA (blue) and *olig2:*EGFP (green) expression. Developmental stages noted at the top. (B) UMAP visualization of the scRNA-seq dataset from *olig2:*EGFP^+^ spinal cord cells obtained from 24, 36 and 48 hpf *Tg(olig2:EGFP)* embryos. Each point represents one cell (n=6489). Colors represent sample time points. (C-F) UMAP plots of selected transcripts. Cells are colored by expression level (gray is low, purple is high). *prdm8* expression overlaps extensively with *olig2, sox19a* and *nkx2*.*2a*. (G-J) Representative transverse trunk spinal cord sections processed for fluorescent ISH to detect *olig2, prdm8* and *nkx2*.*2a* mRNA. (G) 24 hpf (H) 36 hpf (I) 48 hpf (J) 72 hpf; arrowheads indicate *prdm8*^+^*/nkx2*.*2a*^+^*/olig2*^+^ cells; solid arrows denote *prdm8*^+^*/nkx2*.*2a*^*–*^*/olig2*^+^ cells; dashed arrows label *prdm8*^*–*^*/nkx2*.*2a*^+^*/olig2*^+^ cells; dashed ovals outline the spinal cord. Scale bars: 10 μm.

Following motor neuron formation, pMN progenitors begin to form OPCs. This is initiated by the reorganization of ventral progenitor domains, such that pMN cells that enter the oligodendrocyte lineage begin to co-express Nkx2.2 and Olig2 (Agius et al., 2004; Fu et al., 2002; Kessaris et al., 2001; Soula et al., 2001; Zhou et al., 2001). Our scRNA-seq data show that at 48 hpf, cells that expressed *nkx2*.*2a* and *olig2* also expressed *prdm8*, signifying that nascent OPCs express *prdm8* (Fig 1F). We validated these observations using fluorescent RNA ISH. Consistent with our scRNA-seq data, *prdm8* mRNA puncta were present in the pMN domain marked by *olig2* expression at 24, 36, and 48 hpf (Fig. 1H-J). At 24 and 36 hpf, cells that expressed *prdm8* and *olig2* were adjacent to more ventral *nkx2*.*2a*^+^ p3 domain cells (Fig. 1G-H), but at 48 hpf some cells at the p3/pMN border expressed all three transcripts (Fig. 1I). By 72 hpf, most *olig2* mRNA expression was depleted from the pMN domain but evident at high level in oligodendrocyte lineage cells. At this stage some *olig2*^+^ cells expressed both *prdm8* and *nkx2*.*2a*, some expressed only *nkx2*.*2a* and others expressed only *prdm8* (Fig. 1J). These data indicate that following pMN progenitor cell expression, *prdm8* expression is differentially maintained by oligodendrocyte lineage cells.

To determine the identity of *prdm8*^+^ cells, we compared *prdm8* expression with expression of genes characteristic of the oligodendrocyte lineage in the scRNA-seq dataset. At 48 hpf, a subset of cells that expressed *prdm8* also expressed *sox10*, which encodes a transcription factor expressed by all oligodendrocyte lineage cells (Britsch et al., 2001; Kuhlbrodt et al., 1998; Park et al., 2002) (Fig. 2A,B). Some *prdm8*^+^ *sox10*^+^ cells also expressed *myrf*, which encodes Myelin Regulatory Factor, a transcription factor required for oligodendrocyte differentiation (Emery et al., 2009) (Fig. 2C). Our data set includes only a few *mbpa*^+^ cells, and these appeared as a small subset of *sox10*^+^ *myrf*^+^ cells (Fig. 2D). Therefore, these cells represent pre-myelinating oligodendrocytes. A heatmap representation of these cells (Fig. 2E,F) showed that most *sox10*^+^ *nkx2*.*2a*^+^ cells expressed *prdm8* at high levels. However, cells that also expressed *myrf* and *mbpa* at higher levels had little *prdm8* expression. These data suggest that pre-myelinating oligodendrocytes downregulate *prdm8* expression as they differentiate.

**Fig. 2.**
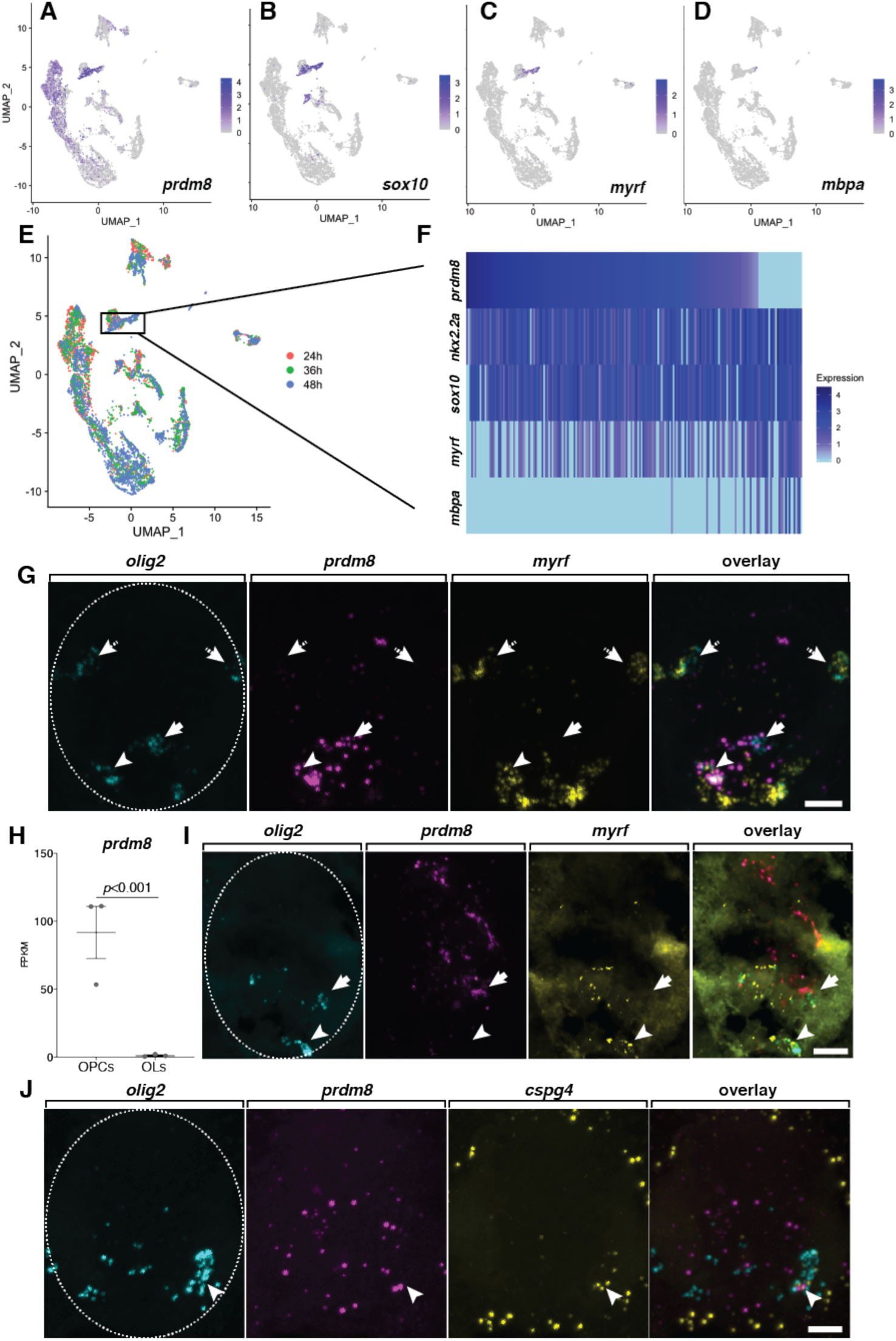
Differentiating oligodendrocytes downregulate *prdm8* expression. (A-D) UMAP plots of select transcripts. Cells are colored by expression level (gray is low, purple is high). *prdm8* expression overlaps considerably with *sox10* and *myrf* but not with *mbpa*. (E) UMAP visualization of subclustered 48 hpf pre-myelinating oligodendrocytes (n=220), indicated by the box. (F) Heatmap showing select transcripts expressed by pre-myelinating oligodendrocytes. (G) Representative transverse trunk spinal cord sections obtained from 72 hpf larvae processed for fluorescent ISH. Dorsal is up. Arrowheads mark *prdm8*^+^*/myrf*^+^*/olig2*^+^ pre-myelinating oligodendrocytes that express *prdm8*; dashed arrows point to *prdm8*^*–*^*/myrf*^+^*/olig2*^+^ pre-myelinating oligodendrocytes that do not express *prdm8*; solid arrows denote *prdm8*^+^*/myrf*^*–*^*/olig2*^+^ OPCs. (H) *prdm8* expression (FPKM) in OPCs (n=3) and oligodendrocytes (n=3) isolated from batched 7 dpf larvae. Data represent mean ± s.e.m. Statistical significance assessed by one-way ANOVA. (I,J) Representative transverse trunk spinal cord sections obtained from 7 dpf larvae processed for fluorescent ISH, dorsal on top. (I) Arrowhead denotes a *prdm8*^*–*^*/myrf*^+^*/olig2*^+^ myelinating oligodendrocyte; solid arrow denotes a *prdm8*^+^*/myrf*^*–*^*/olig2*^+^ OPC. (J) Arrowhead labels a *prdm8*^+^*/cspg4*^+^*/olig2*^+^ OPC; dashed oval outlines the spinal cord boundary. Scale bar: 10 μm.

To corroborate these observations, we compared *prdm8, myrf* and *olig2* mRNA expression using fluorescent RNA ISH in the trunk spinal cord of 72 hpf larvae. Consistent with our scRNA-seq findings, this revealed that a majority of *olig2*^+^ cells expressed either *prdm8* or *myrf* and that few *olig2*^+^ cells expressed both genes (Fig. 2G). To determine if some oligodendrocyte lineage cells maintain *prdm8* expression, we assessed RNA-seq data collected from *cspg4*^+^ OPCs and *mbpa*^+^ oligodendrocytes isolated from 7 days post-fertilization (dpf) larvae (Ravanelli et al., 2018). We found that OPCs expressed *prdm8* at 75-fold higher levels than oligodendrocytes (Fig. 2H). We validated these data using fluorescent RNA ISH to label *prdm8* mRNA at 7 dpf in combination with *myrf* to mark oligodendrocytes (Fig. 2I) or *cspg4* to mark OPCs (Fig. 2J). This revealed that OPCs but not oligodendrocytes expressed *prdm8*, confirming our RNA-seq data. We therefore conclude that pMN progenitors and OPCs express *prdm8* and that *prdm8* expression declines in oligodendrocytes undergoing differentiation.

### Zebrafish larvae lacking *prmd8* function have excess oligodendrocytes and a deficit of OPCs

Zebrafish *prdm8* encodes a 502 amino acid protein containing an N-terminal PR/SET domain and three zinc finger domains, similar to its human and mouse orthologs (Fig. 3A). To investigate Prdm8 function we used CRISPR/Cas9 to create gene-disrupting mutations within the first exon (Fig. 3B). We verified the efficiency of sgRNA targeting using diagnostic fluorescent PCR and subsequently raised injected embryos to adulthood. We identified F0 adults that transmitted mutations through the germ line and selected two, *prdm8*^*co49*^ and *prdm8*^*co51*^, for further analysis. DNA sequencing revealed that the *co49* allele contains a 5 bp insertion whereas the *co51* allele has a 4 bp deletion. Both alleles are predicted to result in translation frameshifting leading to premature translation termination and proteins truncated within the PR/SET domain (Fig. 3B). Because the C-terminal zinc finger domains of mouse Prdm8 are necessary for nuclear localization (Eom et al., 2009), we predict that truncated proteins produced by the *co49* an *co51* alleles are non-functional. Genotyping assays revealed that F1 heterozygous adults transmitted mutant alleles to progeny with Mendelian frequencies (Fig. 3C). Homozygous mutant embryos have no discernable morphological phenotype at 24 hpf (Fig. 3D) or at early larval stages (data not shown).

**Fig. 3.**
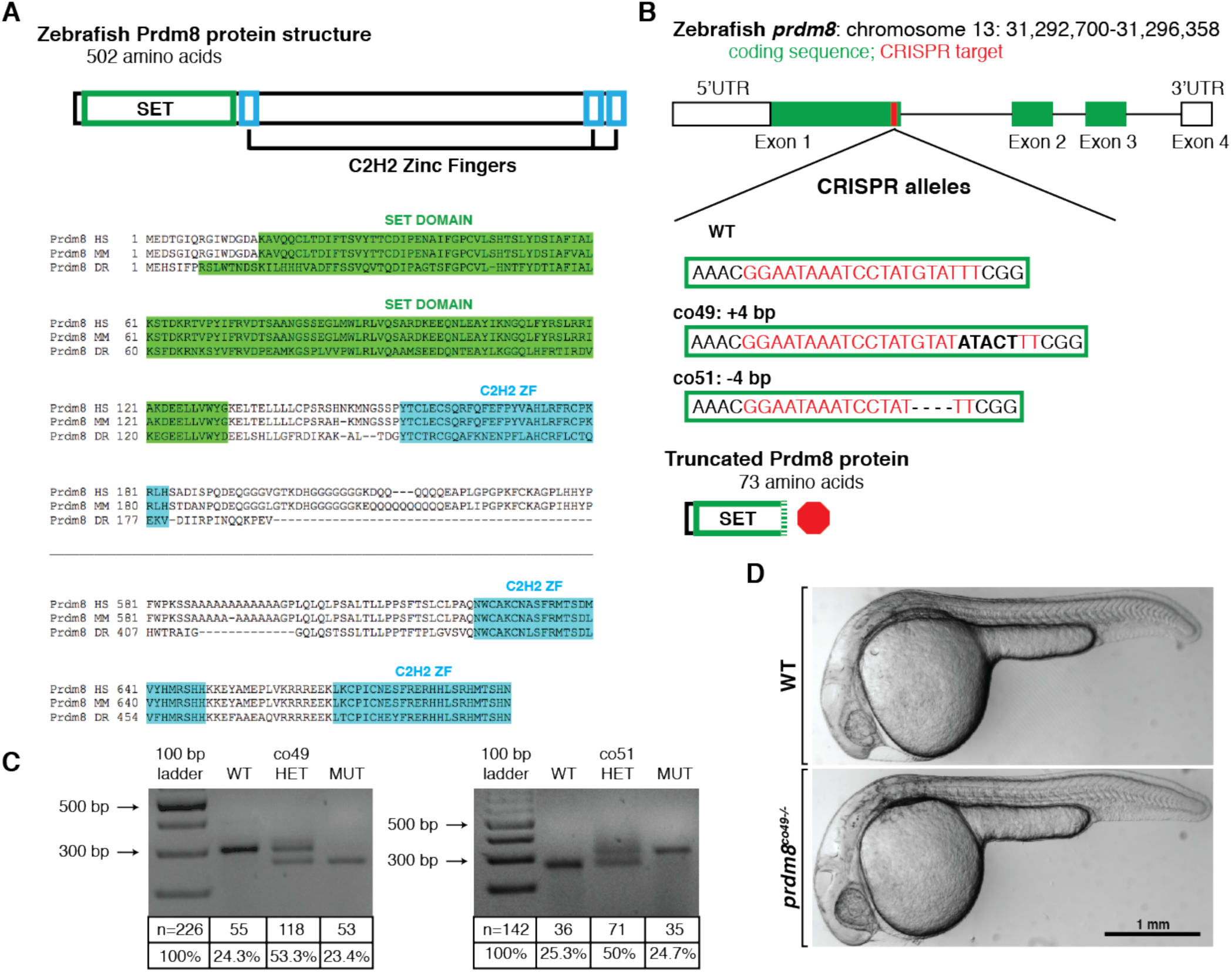
Generation and characterization of *prdm8* loss-of-function mutations. (A) Zebrafish Prdm8 protein structure is depicted as an empty black box with the SET domain highlighted in green and the C2H2 zinc finger domains in blue. Alignment of Prdm8 amino acid sequences from human (HS), mouse (MM), and zebrafish (DR). Conserved SET domain and C2H2 zinc finger domains are shown as green or blue boxes, respectively. (B) Schematic representing *prdm8* gene structure. The sequence targeted for CRISPR/Cas9-mediated mutagenesis is marked by a red line in exon 1. The wild-type sequence CRISPR target sequence is shown below as red text and the *co49* insertion and *co51* deletion are shown as bolded text or dashes, respectively. Both mutations are predicted to produce 73 amino acid proteins truncated at the C-terminal end of the SET domain. (C) Images showing *prdm8* DNA fragmentation following dCAPS genotyping of homozygous wild-type, heterozygous and homozygous mutant embryos with sample genotype frequencies. (D) Representative images of living 24 hpf wild-type and *prdm8*^*co49-/-*^ embryos

To determine if *prdm8* regulates formation of oligodendrocytes, we performed RNA ISH to detect *myrf* expressed by wild-type and *prdm8* mutant larvae. At 72 hpf, larvae homozygous for the *co49* allele had almost twice as many spinal cord *myrf*^+^ oligodendrocytes compared to wild-type siblings (Fig. 4A,B). Heterozygous siblings were not different from wild type (Fig. 4A,B). Larvae homozygous for the *co51* allele similarly had excess *myrf*^+^ oligodendrocytes relative to wild-type siblings (Fig. 4C,D). Larvae trans-heterozygous for the *co49* and *co51* alleles also had a greater number of *myrf*^+^ oligodendrocytes (Fig. 4E,F), indicating that this phenotype results specifically from loss of *prdm8* function and not as a consequence of an off-target mutation produced by CRISPR/Cas9. We additionally examined expression of *mbpa*. Consistent with our *myrf* data, *co49* homozygous mutant larvae had approximately two-fold more dorsal *mbpa*^+^ oligodendrocytes than wild-type siblings (Fig. 4G,H). These data indicate that Prdm8 limits oligodendrocyte formation.

**Fig. 4.**
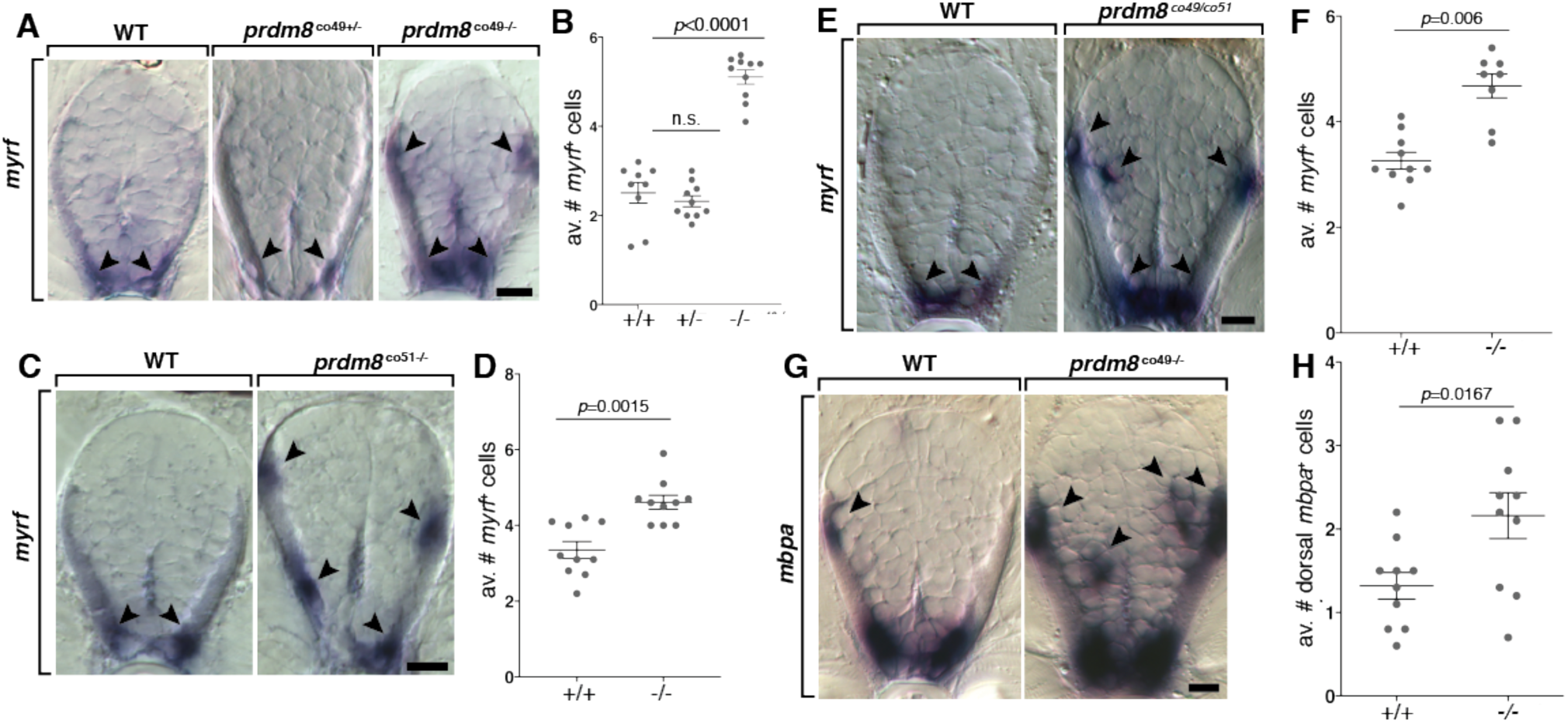
*prdm8* mutant larvae have excess oligodendrocytes. (A,C,E,G) Representative trunk spinal cord transverse sections obtained from 72 hpf larvae showing mRNA expression patterns detected by ISH. Images and quantification of *myrf* expression in wild-type, heterozygous and homozygous *co49* mutant larvae (A,B), wild-type and homozygous *co51* mutant larvae (C,D) and wild-type and *co49/co51* mutant larvae (E,F). Arrowheads mark *myrf*^+^ oligodendrocytes. (G,H) Images of *mbpa* expression and quantification of dorsal *mbpa*^+^ oligodendrocytes in wild-type and homozygous *co49* mutant larvae. Arrowheads denote *mbpa*^+^ oligodendrocytes. n = 10 larvae for each genotype except wild type in B (n = 9), and *co49/co51* mutant larvae in F (n = 8). Data represent mean ± s.e.m. Statistical significance evaluated by Mann-Whitney U test (B, D, F) and Student’s t test (H). Scale bars: 10 μm.

To identify the source of excess oligodendrocytes, we first counted the number of oligodendrocyte lineage cells using an antibody to detect expression of Sox10 in spinal cord sections of wild-type and mutant larvae carrying the *Tg(olig2:EGFP)* reporter. At 72 hpf, homozygous *prdm8* mutant larvae had the same number of Sox10^+^ *olig2*:EGFP^+^ cells as control larvae (Fig. 5A,B). To determine the proportion of Sox10^+^ cells that differentiated as oligodendrocytes, we then performed immunohistochemistry to detect Sox10 on sections obtained from 5 dpf larvae carrying a *Tg(mbpa:tagRFPt)* transgenic reporter. This experiment showed that homozygous *prdm8* mutant larvae had a significant increase in the number of Sox10^+^ *mbpa:*tagRFPt^+^ oligodendrocytes without a change in the total number of Sox10^+^ cells relative to sibling controls (Fig. 5C,D). To assess the OPC population we labeled sections from 5 dpf larvae carrying a *Tg(cspg4:mCherry)* transgenic reporter (Ravanelli et al., 2018) with Sox10 antibody. Homozygous *prdm8* mutant larvae had fewer OPCs than wild-type siblings, but total oligodendrocyte lineage cells were unchanged (Fig. 5E,F). These data indicate that Prdm8 regulates the proportion of OPCs that differentiate as oligodendrocytes.

**Figure 5.**
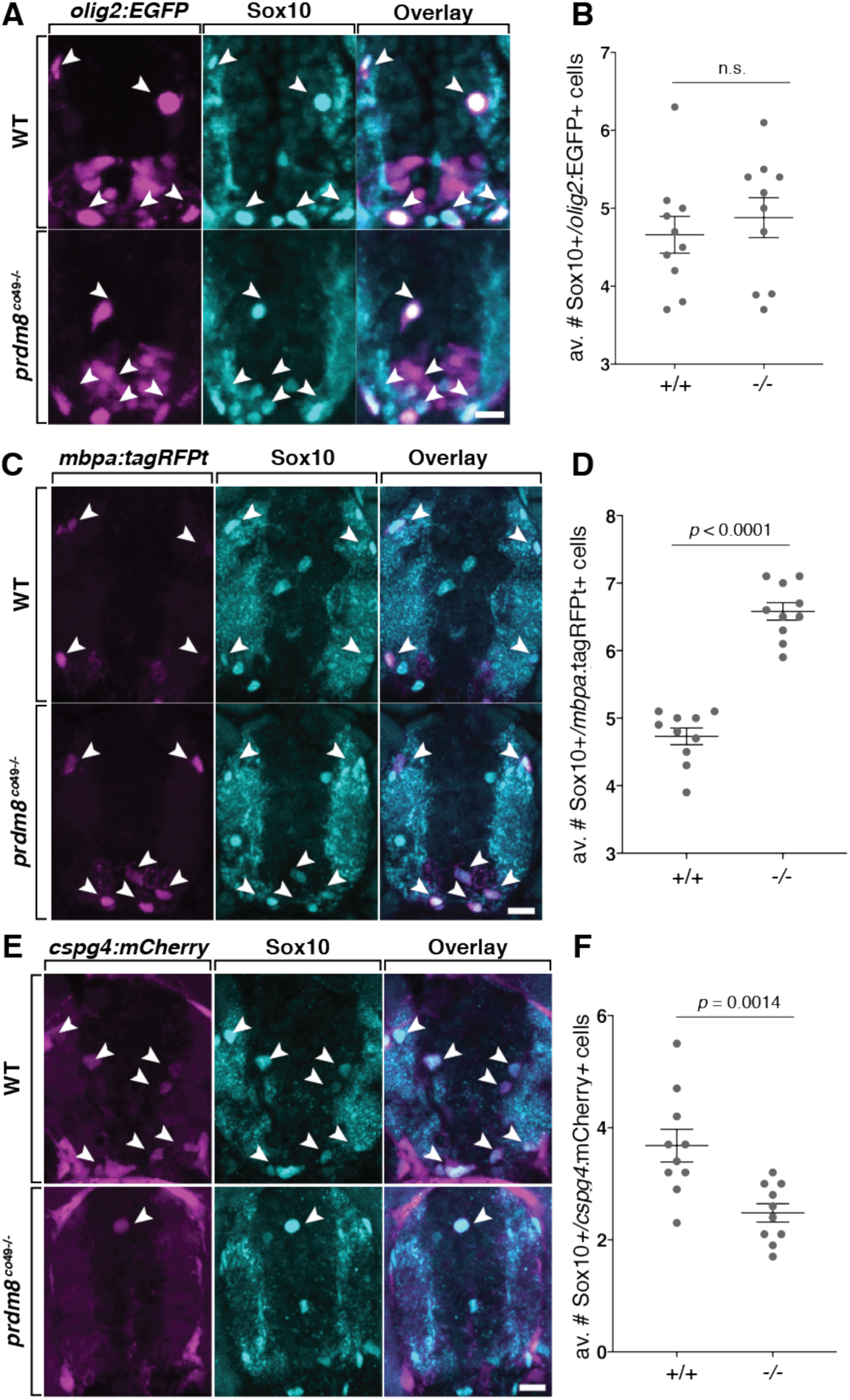
*prdm8* mutant larvae have more myelinating oligodendrocytes and a deficit of OPCs. (A,C,E) Representative images of trunk spinal cord transverse sections processed to detect Sox10 expression (blue) in combination with transgenic reporter gene expression (pink). (A,B) The number of Sox10^+^ *olig2*:EGFP^+^ oligodendrocyte lineage cells is similar in 72 hpf wild-type and *prdm8*^*co49*^ mutant larvae. Arrowheads indicate oligodendrocyte lineage cells. n = 10 for both genotypes. (C,D) 5 dpf *prdm8*^*co49*^ mutant larvae have more Sox10^+^ *mbpa*:tagRFP^+^ oligodendrocytes (arrowheads) than wild-type larvae but there was no difference in total Sox10^+^ oligodendrocyte lineage cells (*prdm8*^*co49*^: 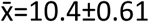, wild-type: 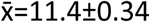, *p*=0.19, graph not depicted). n = 14 for both genotypes. (E,F) 5 dpf *prdm8*^*co49*^ mutant larvae 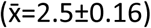 have fewer Sox10^+^ *cspg4*:mCherry^+^ OPCs (arrowheads) than wild-type larvae 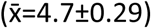, but the number of total Sox10^+^ oligodendrocyte lineage cells was unchanged (*prdm8*^*co49*^: 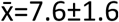, wild-type: 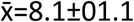, *p*=0.70, graph not depicted). n = 14 for both genotypes. Data represent mean ± s.e.m. Statistical significance evaluated by Student’s *t* test (B, D) and Mann-Whitney U test (F). Scale bars: 10 μm.

### *prdm8* regulates the timing of a neuron-glia switch by repressing neural tube Shh signaling activity

We next investigated whether *prdm8* regulates motor neuron formation from pMN progenitors, which precedes OPC specification. To do so, we used an antibody to detect Isl1/2 (Isl), which marks post-mitotic motor neurons (Ericson et al., 1992). At 24 hpf, homozygous *prdm8* mutant embryos had the same number of Isl^+^ *olig2*:EGFP^+^ motor neurons as controls (Fig. 6A,B), signifying that *prdm8* mutant embryos initiate motor neuron formation normally. By contrast, at 36 and 48 hpf *prdm8* mutant embryos had fewer motor neurons than control embryos (Fig. 6C-F), suggesting that Prdm8 is required to maintain motor neuron production from pMN progenitors.

**Figure 6.**
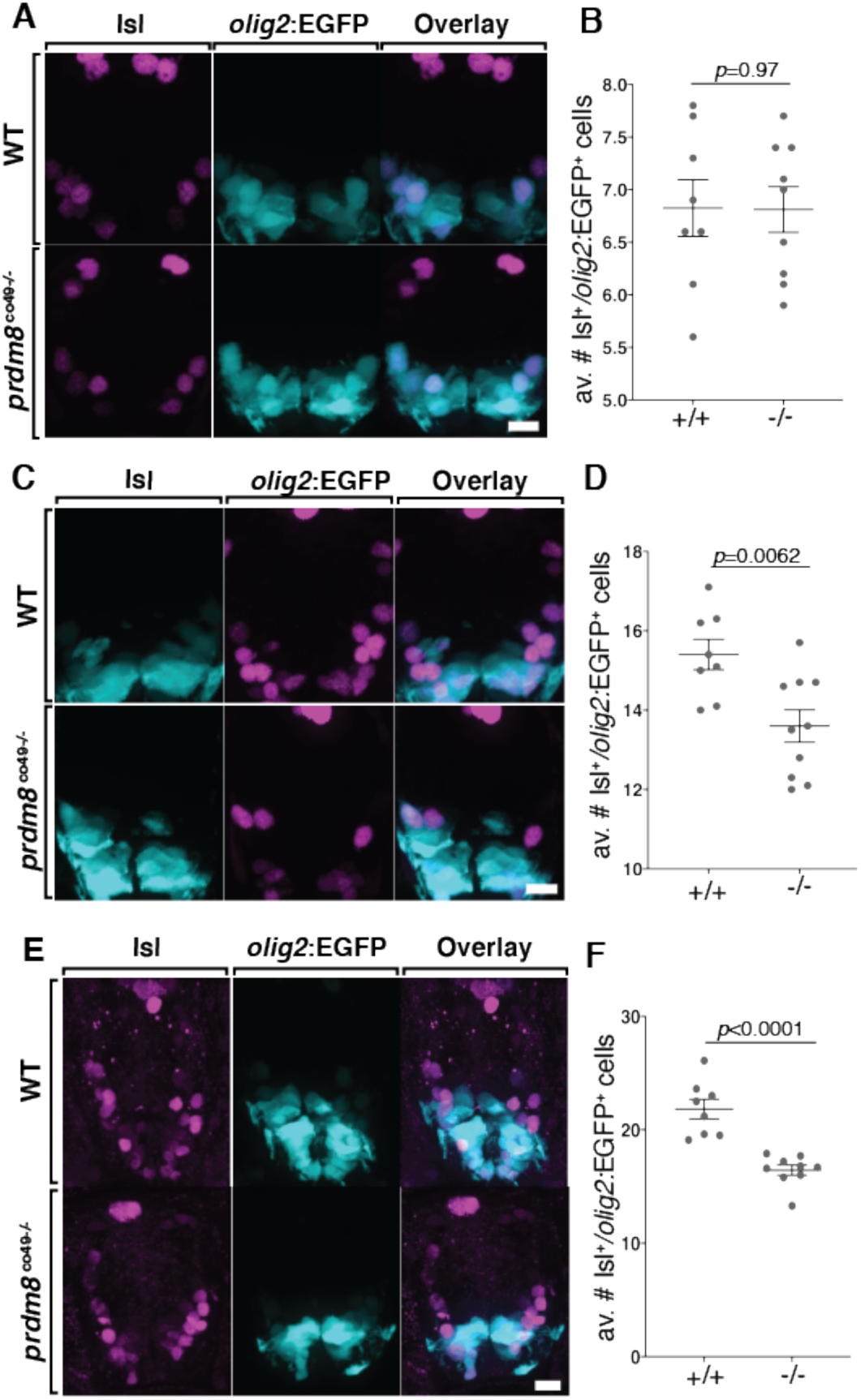
*prdm8* mutant embryos have a deficit of motor neurons. (A,C,E) Representative images of trunk spinal cord transverse sections processed to detect Isl expression (pink) in combination with *olig2:*EGFP (blue). (A,B) The number of Isl^+^ *olig2:*EGFP^+^ motor neurons is similar in 24 hpf wild-type (n=8) and *prdm8*^*co49*^ mutant embryos (n=9). (C,D) 36 hpf *prdm8*^*co49*^ mutant embryos (n=10) have fewer Isl^+^ *olig2:*EGFP^+^ motor neurons than wild-type embryos (n=8). (E,F) 48 hpf *prdm8*^*co49*^ mutant embryos (n=9, 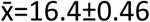) have fewer Isl^+^ *olig2:*EGFP^+^ motor neurons than wild-type embryos (n=8, 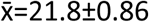). Data represent mean ± s.e.m. Statistical significance evaluated by Student’s *t* test (B,F) and Mann-Whitney U test (D). Scale bars: 10 μm.

One possible explanation for these data is that in the absence of *prdm8* function, pMN progenitors prematurely switch from motor neuron to OPC production, resulting in a deficit of motor neurons. To test this prediction, we exposed 24, 30 and 36 hpf embryos to a pulse of BrdU to label cells in S-phase and used immunohistochemistry at 48 hpf to identify the progeny of the labeled progenitors (Fig. 7A). Compared to stage-matched wild-type control embryos, homozygous *prdm8* mutant embryos exposed to BrdU at each time point had a deficit of BrdU^+^ Isl^+^ motor neurons (Fig. 7B-H). By contrast, mutant embryos pulsed with BrdU at 24 and 30 hpf had more BrdU^+^ Sox10^+^ cells than controls (Fig. 7B,C,E,F,I) whereas those pulsed at 36 hpf had fewer (Fig. 7D,G,I). These data indicate that *prdm8* mutant embryos prematurely terminate motor neuron formation and concomitantly produce OPCs earlier than normal. However, mutant embryos also prematurely terminate OPC production. Altogether these data raise the possibility that Prdm8 prevents premature OPC specification, thus preserving pMN progenitors for motor neuron fate.

**Figure 7.**
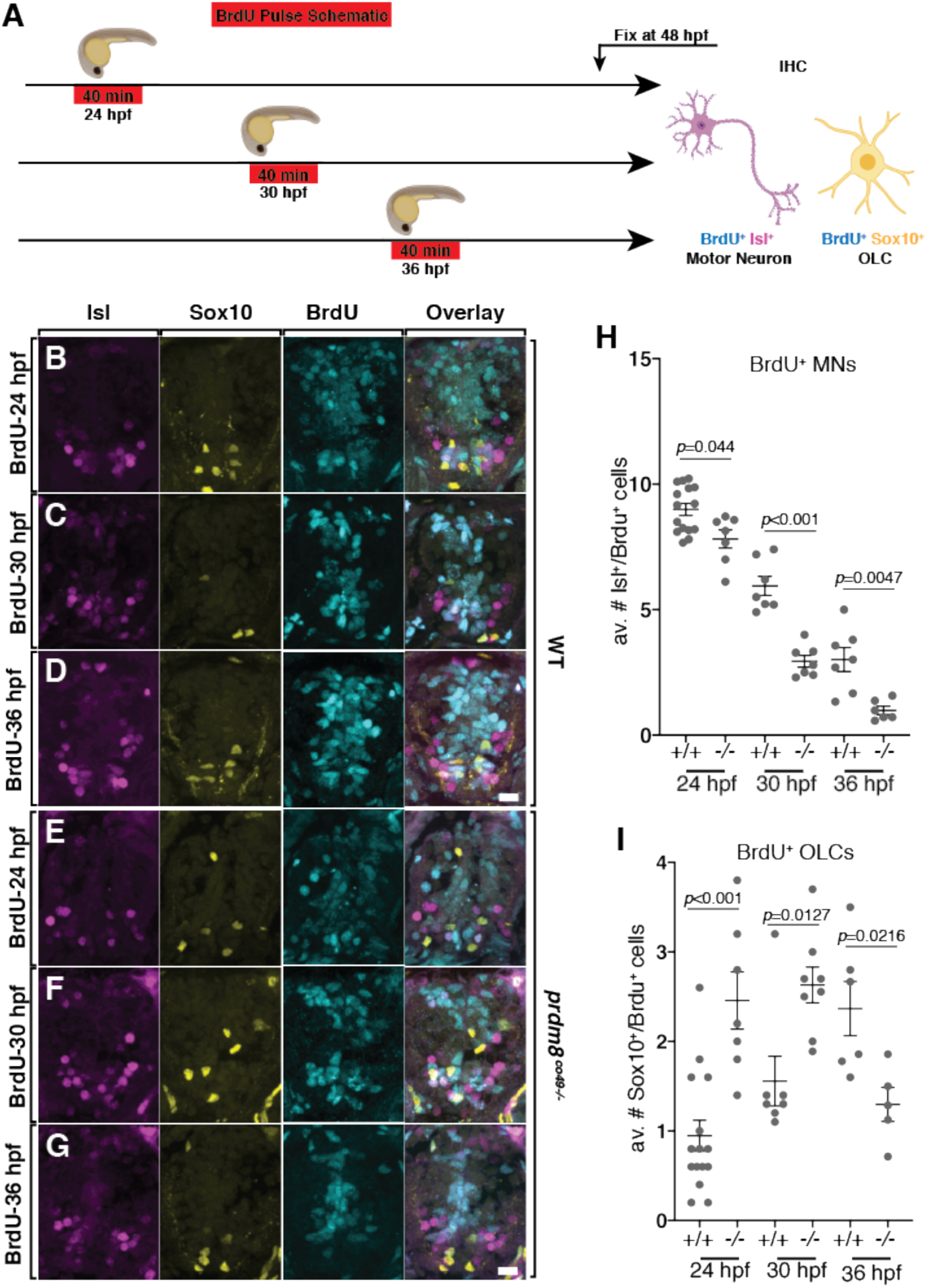
*prdm8* mutant embryos prematurely switch from motor neuron to OPC production. (A) Schematic of BrdU pulses. (B-G) Representative images of trunk spinal cord sections from 48 hpf embryos treated with BrdU and processed to detect Isl (pink), Sox10 (yellow) and BrdU (blue). Wild-type (B) and *prdm8*^*co49-/-*^ (E) embryos pulsed with BrdU at 24 hpf. Wild-type (C) and *prdm8*^*co49-/-*^ (F) embryos pulsed with BrdU at 30 hpf. Wild-type (D) and *prdm8*^*co49-/-*^ (G) embryos pulsed with BrdU at 36 hpf. (H) Quantification of Isl^+^/BrdU^+^ motor neurons pulsed with BrdU at 24 hpf in wild-type (n=15) and *prdm8*^*co49-/-*^ (n=7); 30 hpf in wild-type (n=7) and *prdm8*^*co49-/-*^ (n=7,); 36 hpf in wild-type (n=7) and *prdm8*^*co49-/-*^ (n=6). (I) Quantification of Sox10^+^/BrdU^+^ cells pulsed with BrdU at 24 hpf in wild-type (n=15) and *prdm8*^*co49-/-*^ (n=7); 30 hpf in wild-type (n=7) and *prdm8*^*co49-/-*^ (n=8); 36 hpf in wild-type (n=6) and *prdm8*^*co49-/-*^ (n=5). Data represent mean ± s.e.m. Statistical significance evaluated by Manny-Whitney U test. Analysis of embryos pulsed with BrdU at 24 hpf represent data collected from two laboratory replicates. Scale bars: 10 μm.

OPC specification coincides with and requires a dorsal expansion of Nkx2.2 expression from the p3 domain, resulting in co-expression of Nkx2.2 and Olig2 (Kessaris et al., 2001; Kucenas et al., 2008; Soula et al., 2001; Xu et al., 2000). Thus, one possible mechanism by which *prdm8* prevents premature OPC specification is by regulating the time at which pMN progenitors express *nkx2*.*2a*. To examine this possibility, we used fluorescent RNA ISH to label *nkx2*.*2a* and *olig2* mRNA and quantified *nkx2*.*2a* puncta in the pMN domain at 28 hpf, prior to OPC specification (Fig. 8A-B). *prdm8* mutant embryos had more *nkx2*.*2a* mRNA puncta localized to pMN cells than controls (Fig. 8A-B). Next, we examined the number of *olig2:*EGFP^+^ OPCs that express *nkx2*.*2a* at 48 hpf. *prdm8* mutant embryos had almost 3-fold more dorsal *nkx2*.*2a*^+^ *olig2:*EGFP^+^ OPCs compared to controls (Fig. 8C-D). Together, these data suggest that *prdm8* controls the timing of OPC specification by controlling the time at which pMN cells initiate *nkx2*.*2a* expression.

**Figure 8.**
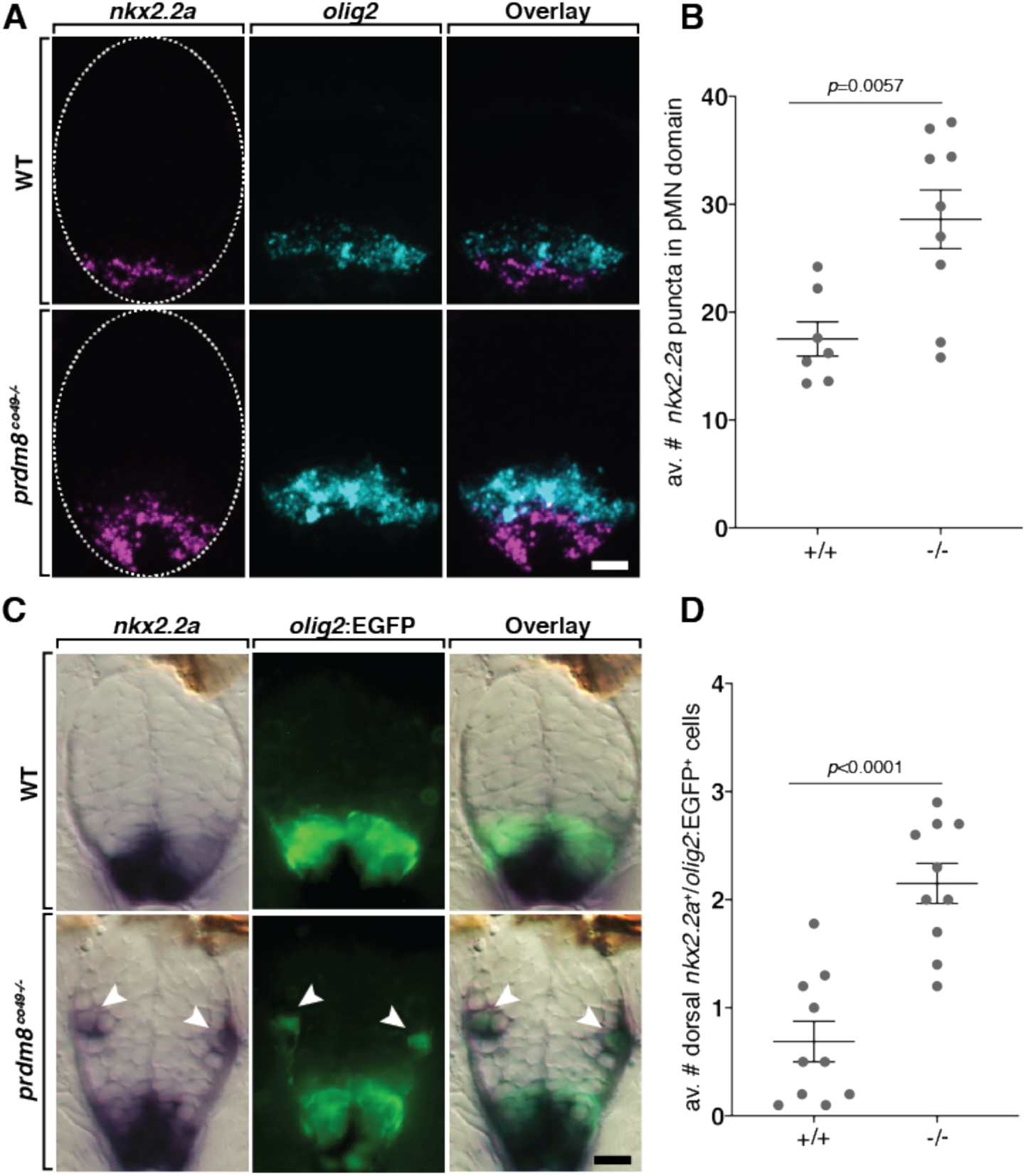
pMN cells prematurely express *nkx2.2a* in *prdm8* mutant embryos. (A) Representative transverse trunk spinal cord sections obtained from 28 hpf embryos processed for fluorescent ISH to detect *olig2* (blue) and *nkx2.2a* (pink) mRNA. (B) More *nkx2.2a* puncta are located within the *olig2*^+^ pMN domain of *prdm8*^*co49-/-*^ embryos (n=9) compared to wild-type embryos (n=7). (C) Representative transverse sections of trunk spinal cord obtained from 48 hpf embryos showing *nkx2.2a* RNA (blue) and *olig2:*EGFP (green) expression. Arrowheads indicate dorsally migrated OLCs. (D) *prdm8*^*co49-/-*^ (n=10) have more dorsal OPCs (*nkx2.2a*^+^*/olig2*:EGFP^+^) than wild-type embryos (n=10) at 48 hpf. Data represent mean ± s.e.m. Statistical significance evaluated by Student’s *t* test. Scale bars: 10 μm.

At the end of neurogenesis, ventral spinal cord cells transiently elevate Shh signaling activity, which is necessary for OPC specification (Orentas et al., 1999; Soula et al., 2001; Touahri et al., 2012). Experimentally increasing Shh levels caused premature termination of motor neuron formation and precocious OPC formation (Danesin et al., 2006), similar to the loss of *prdm8* function and thereby raising the possibility that Prdm8 suppresses Shh activity in the ventral spinal cord. To test this possibility, we probed for expression of *ptch2*, a transcriptional target of the Shh signaling pathway. At 24 hpf *prdm8* mutant embryos appeared to express more *ptch2* than wild-type embryos (Fig. 9A). By 48 hpf there was no visible difference in *ptch2* expression between genotypes (Fig. 9A). Next we used fluorescent RNA ISH to quantify *ptch2* expression. At 24 and 36 hpf *prdm8* mutant embryos expressed more *ptch2* mRNA relative to total spinal cord area compared to controls (Fig. 9B-D). Wild-type and *prdm8* mutant embryos expressed *shha* similarly, suggesting that the elevated level of *ptch2* expression results from increased Shh signaling activity is independent of ligand expression (Fig. 9E). These results are consistent with the possibly that Prdm8 suppresses Shh response in pMN cells to regulate the transition between motor neuron and OPC formation.

**Figure 9.**
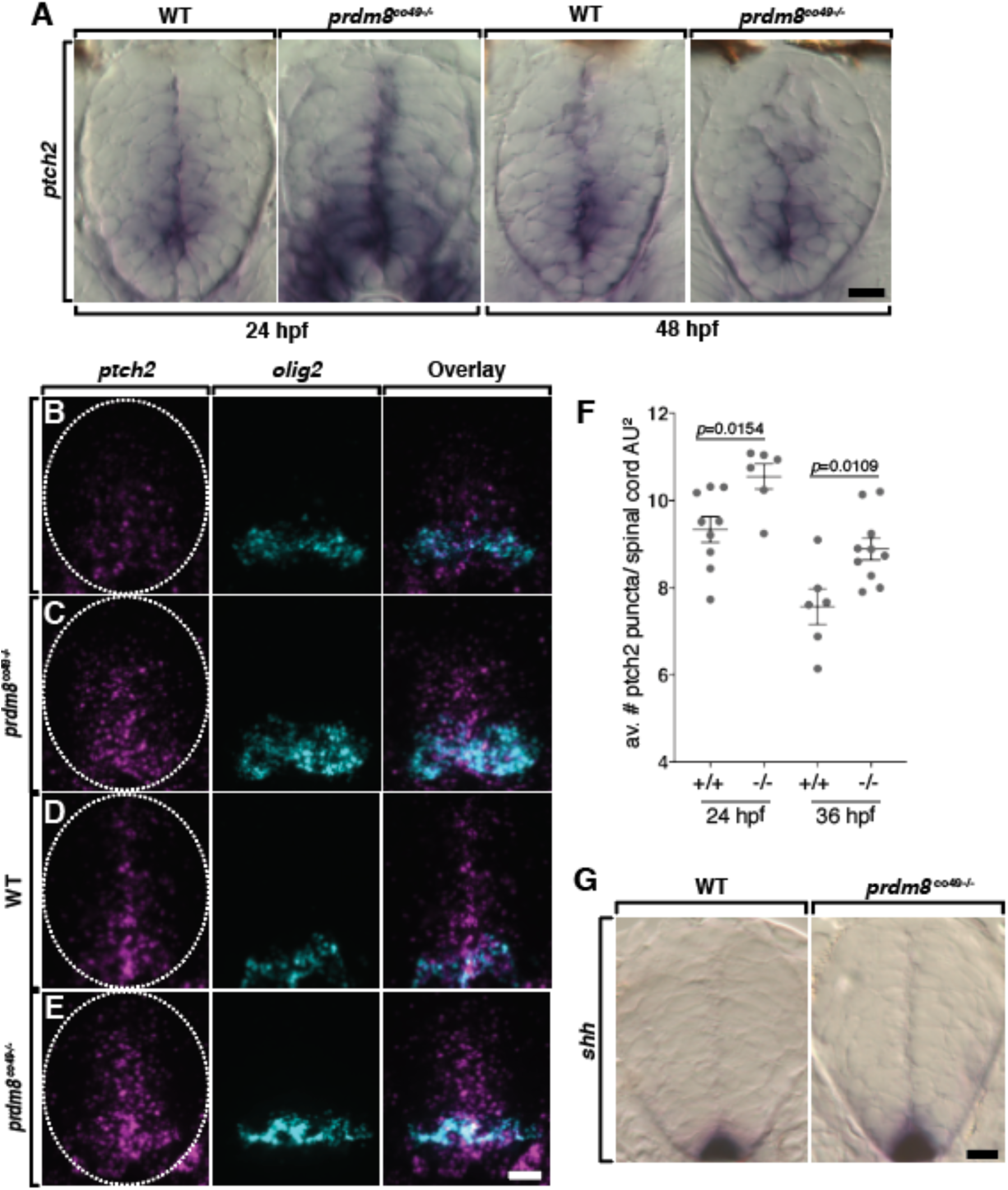
Spinal cord cells of *prdm8* mutant embryos have elevated Shh signaling activity. (A) Representative transverse sections of trunk spinal cords obtained from 24 and 48 hpf wild-type and *prdm8*^*co49-/-*^ embryos, with dorsal up, showing *ptch2* RNA expression. (B-C) Representative transverse trunk spinal cord sections processed for fluorescent ISH to detect *olig2* (blue) and *ptch2* (pink) mRNA at 24 hpf (B) and 36 hpf (C). (D) *prdm8*^*co49-/-*^ embryos have more *ptch2* puncta per AU^2^ of spinal cord at 24 hpf (n=6) and 36 hpf (n=10) than wild-type embryos at 24 hpf (n=9) and 36 hpf (n=6). (E) Representative transverse sections of trunk spinal cord, with dorsal up, showing *shha* RNA expression in 24 hpf wild-type and *prdm8*^*co49-/-*^ embryos. Data represent mean ± s.e.m. Statistical significance evaluated by Student’s *t* test. Dashed oval outlines the spinal cord boundary. Scale bars: 10 μm.

Because the premature transition between motor neuron and OPC production resulting from lack of *prdm8* function resembled the effect of abnormally elevated Shh signaling, we predicted that the number of motor neurons in *prdm8* mutant embryos could be rescued by inhibiting Shh activity. To test this prediction, we treated *prdm8* mutant and wild-type embryos with a low concentration of cyclopamine to partially block Shh signal transduction from 18-30 hpf and assessed motor neuron number at 48 hpf. The number of motor neurons was similar in *prdm8* mutant embryos treated with cyclopamine and wild-type embryos treated with vehicle control (Fig. 10A,B), consistent with our prediction. Furthermore, both wild-type and *prdm8* mutant embryos treated with cyclopamine had more motor neurons than their genotype-matched controls (Fig. 10B), raising the possibility that suppression of Shh signaling delays the motor neuron to OPC switch, resulting in formation of excess motor neurons. By contrast, treatment with cyclopamine from 30-42 hpf, after most spinal cord neurogenesis is normally completed, had no effect on motor neuron number in either wild-type or *prdm8* mutant embryos (Fig. 10C,D). Consistent with our previous assessments, vehicle control-treated mutant embryos had fewer motor neurons than wild-type siblings (Fig. 10A-D). These data therefore support the possibility that Prdm8 suppresses Shh signaling within pMN cells to regulate the termination of motor neuron production and the timing of the neuron-glia switch.

**Figure 10.**
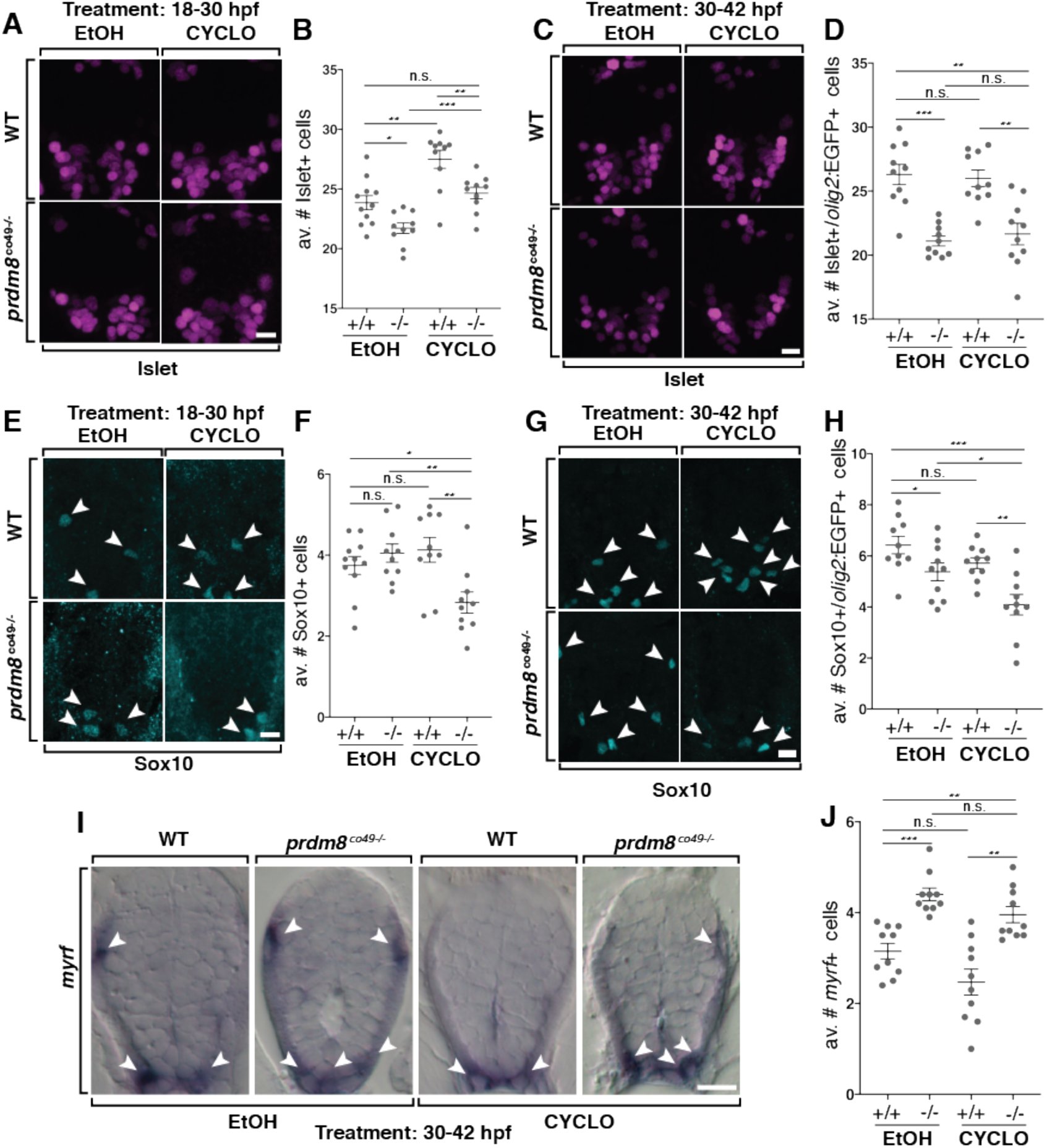
Shh inhibition rescues the motor neuron but not oligodendrocyte phenotypes of *prdm8* mutant embryos. (A,C,E,G) Representative images of trunk spinal cord sections from 48 hpf embryos treated with 0.5 μM cyclopamine (CYCLO) or ethanol (EtOH) from 18-30 hpf (A,E) or 30-42 hpf (C, G) and processed to detect Isl (A,C) or Sox10 (E,G) expression. (A,B) Wild-type embryos treated with EtOH control solution and *prmd8* mutant embryos treated with cyclopamine have similar numbers of motor neurons. (C,D) There are fewer motor neurons (Islet^+^) in *prdm8*^*co49-/-*^ embryos treated with EtOH and cyclopamine compared to wild-type embryos treated with EtOH. (E,F) There are fewer OPCs (Sox10^+^;arrowheads) in *prdm8*^*co49-/-*^ embryos treated with cyclopamine and no difference in OPCs *prdm8*^*co49-/-*^ embryos treated with EtOH compared to wild-type embryos treated with EtOH. (G,H) There are fewer OPCs (Sox10+; arrowheads) in *prdm8*^*co49-/-*^ embryos treated with cyclopamine and slightly less OPCs in *prdm8*^*co49-/-*^ embryos treated with EtOH compared to wild-type embryos treated with EtOH. (I) Representative trunk spinal cord transverse sections obtained from 72 hpf larvae treated with 0.5 μM cyclopamine or ethanol (EtOH) from 30-42 hpf showing *myrf* mRNA expression detected by in situ RNA hybridization. (I,J) *prdm8*^*co49-/-*^ embryos treated with EtOH or cyclopamine have more oligodendrocytes (*myrf*^+^;arrowheads) than wild-type embryos treated with EtOH. n= 10 for all genotypes and treatments expect n=11 for wild-type embryos treated with EtOH (A,E). Data represent mean ± s.e.m. Statistical significance evaluated by Student’s *t* test. \**p*<0.05, \*\**p*<0.001, \*\*\**p*<0.0001. Scale bars: 10 μm.

We next tested whether suppressing Shh signaling in *prdm8* mutant embryos affects oligodendrocyte development. As above, the number of Sox10^+^ cells was not different between wild-type and mutant larvae treated with a control solution between 18-30 hpf (Fig. 10E,F). Whereas wild-type embryos treated with control solution or 0.5 μM cyclopamine from 18-30 hpf had similar numbers of Sox10^+^ cells, *prdm8* mutant embryos treated with cyclopamine had a significant deficit of Sox10^+^ cells compared to mutant embryos treated with control solution and wild-type embryos treated with cyclopamine (Fig. 10E,F). Treating embryos with cyclopamine from 30-42 hpf similarly caused *prdm8* mutant embryos to have fewer Sox10^+^ cells than mutant embryos treated with control solution and wild-type embryos treated with cyclopamine (Fig. 10G,H). One possible explanation for these observations is that, although cyclopamine treatment delays the premature neuron-glia transition in *prdm8* mutant embryos it does not rescue the premature termination of OPC production, thereby resulting in a deficit of oligodendrocyte lineage cells.

We showed above that *prdm8* mutant larvae have excess oligodendrocytes and a deficit of *cspg4*^+^ OPCs. To learn if this phenotype results from misregulated Shh signaling, we treated embryos with cyclopamine from 30-42 hpf and examined expression of *myrf* as a marker for myelinating oligodendrocytes. *prdm8* mutant larvae treated with control solution and cyclopamine had similar numbers of *myrf*^+^ oligodendrocytes (Fig. 10I,J). Thus, suppressing Shh signaling did not rescue the excess oligodendrocyte phenotype of *prdm8* mutant larvae, raising the possibility that Prdm8 regulates oligodendrocyte formation independently of its role in controlling the timing of a neuron-glia switch.

## DISCUSSION

The neuron-glia switch, whereby neural progenitors produce neurons followed by glia, is a general feature of developing nervous systems (Rowitch and Kriegstein, 2010). Despite its important role in diversifying neural cell fate, the mechanisms that cause the switch and determine its timing remain poorly understood. In the ventral spinal cord, a temporally regulated rise in Shh signaling activity appears to trigger pMN progenitors to switch from motor neuron to OPC production (Danesin and Soula, 2017). Our results presented here now indicate that Prdm8 suppresses Shh signaling activity within pMN progenitors to control the timing of the motor neuron-OPC switch.

Distinct types of neurons and glia arise from distinct subpopulations of progenitor cells aligned along the dorsorventral axis of the spinal cord. A large body of work conducted over the past 30 years has shown that the identities of these progenitor populations are determined by combinatorial expression of an extensive array of bHLH and homeodomain transcription factors (Sagner and Briscoe, 2019). Additionally, specific subpopulations of spinal cord progenitors also express members of the Prdm protein family, although how these factors contribute to spinal cord development has received considerably less attention (Zannino and Sagerstrom, 2015). For example, dorsal progenitors express Prdm13, which regulates the balance between excitatory and inhibitory interneuron production by blocking the activity of bHLH transcription factors that drive expression of genes required for excitatory neuron differentiation (Chang et al., 2013; Mona et al., 2017). p1 progenitors express Prdm12, which is required for V1 interneuron formation (Thélie et al., 2015) and Prdm14 promotes Islet2 expression and axon outgrowth in motor neurons (Liu et al., 2012). Finally, pMN, p1 and p2 progenitors express Prdm8 (Kinameri et al., 2008; Komai et al., 2009). Although the spinal cord function of Prdm8 had not been previously investigated, evidence indicating that Prdm8 regulates specification of retinal cells (Jung et al., 2015) and that Prdm8 can interact with bHLH transcription factors (Ross et al., 2012; Yildiz et al., 2019) raises the possibility that Prdm8 contributes to mechanisms that determine spinal cord progenitor fate.

The first main finding of our work is that zebrafish spinal cord cells express *prdm8* similarly to mouse (Kinameri et al., 2008; Komai et al., 2009). Our data show that pMN progenitors express *prdm8* throughout developmental neurogenesis and gliogenesis, but that *prdm8* expression subsides from the ventral spinal cord as pMN progenitors are depleted at the onset of the larval period. Furthermore, our analysis extends the mouse expression data by showing that oligodendrocyte lineage cells also express *prdm8*. Specifically, our data indicate that newly specified OPCs express *prdm8* but then downregulate it as they initiate oligodendrocyte differentiation. By contrast, larval OPCs, marked by *cspg4* expression, maintain *prdm8* expression. Together, these observations raise the possibility that Prdm8 regulates pMN progenitor fate specification and, subsequently, OPC differentiation.

Our second main conclusion is that Prdm8 regulates the timing of the motor neuron to OPC switch by determining how strongly pMN progenitors respond to Shh signaling. Specifically, we found that *prdm8* mutant embryos have a deficit of late-born motor neurons because of a premature neuron-glia switch, that mutant spinal cord cells express abnormally high levels of *ptch2* and that the motor neuron deficit was rescued by treating mutant embryos with a low concentration of a Shh inhibitor. This finding supports previous evidence that a transient burst of Shh signaling activity initiates the switch from motor neuron to OPC production (Danesin and Soula, 2017). This burst is mediated, at least in part, by Sulfatase function (Al Oustah et al., 2014; Danesin et al., 2006; Jiang et al., 2017). Sulfatases are secreted by ventral spinal cord cells and increase the range of Shh ligand in the extracellular matrix by regulating the sulfation state of heparan sulfate proteoglycans (Farzan et al., 2008; Yan and Lin, 2009). How cells receive and process extracellular signals also can influence signaling strength. In particular, Notch signaling increases the sensitivity of neural cells to Shh signaling (Kong et al., 2015; Ravanelli et al., 2018; Stasiulewicz et al., 2015). Currently, we do not know how Prdm8 suppresses Shh signaling activity within pMN progenitors. Because Prdm8 functions as a transcriptional inhibitor (Chen et al., 2018; Eom et al., 2009; Iwai et al., 2018; Ross et al., 2012), Prdm8 might suppress expression of factors that transduce Shh signaling. For example, Prdm8 could suppress expression of the Shh co-receptors Boc, Cdon and Gas1, which enhance cell response to Shh (Allen et al., 2007; Allen et al., 2011). Alternatively, Prdm8 could limit expression of Notch signaling effectors that enhance Shh signaling. Identification of genes misregulated in *prdm8* mutant embryos combined with determination of genomic loci targeted by Prdm8 should help uncover the regulatory function of Prdm8 in pMN progenitor specification.

Finally, we found that *prdm8* mutant larvae have excess oligodendrocytes at the apparent expense of OPCs. There are at least two possible explanations for this phenotype. Because our expression data show that cells undergoing oligodendrocyte differentiation downregulate *prdm8* expression, the first possibility is that Prdm8 inhibits OPC differentiation and, therefore, in its absence, OPCs that normally persist into larval stage instead develop as myelinating oligodendrocytes. A second possibility it that Prdm8 regulates allocation of pMN progenitors for distinct oligodendrocyte lineage cell fates. Previously, in a process we called progenitor recruitment, we showed that motor neurons, OPCs that rapidly differentiate and OPCs that persist into larval stage arise from distinct pMN progenitors that sequentially initiate *olig2* expression (Ravanelli and Appel, 2015; Ravanelli et al., 2018). We also found that slightly higher levels of Shh signaling favors formation of oligodendrocytes over larval OPCs, which is similar to the oligodendrocyte phenotype of *prdm8* mutant animals. However, inhibiting Shh with cyclopamine did not restore oligodendrocytes and OPCs to their normal numbers, raising the possibility that Prdm8 regulates oligodendrocyte lineage cell fate independently of Shh signaling. Identifying Prdm8 regulatory targets combined with detailed cell lineage analysis will help us discriminate between these possibilities.

## METHODS AND MATERIALS

### Zebrafish lines and husbandry

All animal work was approved by the Institutional Animal Care and Use Committee (IAUCUC) at the University of Colorado School of Medicine. All non-transgenic embryos were obtained from pairwise crosses of males and females from the AB strain. Embryos were raised at 28.5°C in E3 media (5 mM NaCl, 0.17 mM KCl, 0.33 mM CaCl_2_, 0.33 mM MgSO_4_ at pH 7.4, with sodium bicarbonate), sorted for good health and staged accordingly to developmental morphological features and hours post-fertilization (hpf) (Kimmel et al., 1995). Developmental stages are described in the results section for individual experiments. Sex cannot be determined at embryonic and larval stages. Embryos were randomly assigned to control and experimental conditions for BrdU and pharmacological treatments. The transgenic lines used were *Tg(olig2:EGFP)*^*vu12*^ (Shin et al., 2003), *Tg(mbpa:tagRFPt)*^*co25*^ (Hines et al., 2015) and *Tg(cspg4:mCherry)*^*co28*^ (Ravanelli et al., 2018). All transgenic embryos were obtained from pairwise crosses of males or females from the AB strain to males or females of each transgenic line used.

### Generation of CRISPR/Cas9 mutant zebrafish lines

We designed a single guide RNA (sgRNA) for the zebrafish *prdm8* gene using the CRISPOR web tool (http://crispor.tefor.net/) (Table 1). The sgRNA was constructed by annealing sense and anti-sense single stranded oligonucleotides containing 5’ Bsa1 restriction overhangs and was inserted into Bsa1 linearized pDR274 with the Quick Ligase Kit (NEB) (Table 1). The plasmid was transformed into chemically competent DH5a cells and purified from individual colony liquid cultures with the Qiagen Spin Miniprep Kit (Qiagen). To make the sgRNA we linearized purified pDR274 containing the guide sequence with Dra1 and used a T7 RNA polymerase for in vitro transcription (NEB). The pMLM3613 plasmid encoding *cas9* was used for in vitro transcription using the SP6 mMessage mMachine Kit (Ambion) according to manufacturer’s instructions. The sgRNA and *cas9* mRNA were co-injected into single-cell AB zebrafish embryos at the following concentrations: 200 ng/μl *cas9* mRNA and 150 ng/μl *prdm8* mRNA.

The following day injected embryos were assayed for sgRNA activity by DNA extraction and two rounds of PCR amplification, first to amplify the *prdm8* CRIPSR target with gene specific primers containing a M13F extension to the 5’ end of the forward primer (5’TGTAAAACGACGGCCAGT3’) and a second to add a fluorescein tag to the 5’ end of the amplified region (Table 2). The fluorescein tagged PCR product was analyzed using capillary gel electrophoresis to detect product length. To detect F0 founders, we set up pairwise crosses of injected adults with ABs and screened their offspring for mutagenic events by fluorescent PCR and capillary gel electrophoresis. We used Sanger sequencing to determine the sequence of mutant alleles. We identified two mutant alleles, one with a 5 bp insertion (*prdm8*^*co49*^) and another with a 4 bp deletion (*prdm8*^*co51*^) (Fig. 3B). These lines were maintained as heterozygotes through pairwise crosses with ABs or transgenic lines.

### dCAPS Genotyping

To genotype embryos and adults we designed a derived cleaved amplified polymorphic sequencing (dCAPs) assay to insert a restriction site into either the mutant or wild type allele via PCR. Specific forward primers were designed for each allele, one that added a BsrG1 restriction site into the *prdm8*^*co49*^ allele and another that added a Nde1 restriction site into the wild-type allele for *prdm8*^*co51*^ identification (Table 2). PCR products were digested with appropriate enzymes and samples were run on a 2.5% agarose gel; the *prdm8*^*co49*^ digest creates 267 and 37 bp mutant digested fragments and a 299 bp undigested wild type fragment; the *prdm8*^*co51*^ digest creates two digested wild-type fragments of 260 and 55 bp and a 315 bp undigested mutant fragments (Fig. 3C).

### BrdU Labeling

Embryos and larvae were dechorionated, incubated in a 20 μM BrdU solution for 40 min on ice at indicated time points (Fig. 7A) and subsequently washed 4 × 5 min with embryo medium. Embryos and larvae were allowed to develop until 2 dpf in embryo E3 media. Samples were fixed in 4% paraformaldehyde (PFA) in 1X PBS, embedded (1.5% agar, 5% sucrose), sectioned and prepared for immunohistochemistry as described below.

### Cyclopamine Treatment

Cyclopamine (Cat#11321, Cayman Chemical) was reconstituted in ethanol to make a 10 mM stock and stored at -20°C. Dechorinated embryos were treated with 0.5 μM cyclopamine or an equal concentration of ethanol alone in E3 media at indicated time points. Following treatment, embryos were washed three times with E3 media and grown to designated time points before fixation.

### Whole Mount In situ RNA Hybridization

In situ RNA hybridizations were performed as described previously (Hauptmann and Gerster, 2000). Probes included *ptch2* (Concordet et al., 1996), *nkx2.2a* (Barth and Wilson, 1995), *myrf, mbpa* (Brösamle and Halpern, 2002) and *prdm8* (Table 2). Plasmids were linearized with appropriate restriction enzymes and cRNA preparation was carried out using Roche DIG-labeling reagents and T3, T7 or SP6 RNA polymerases (New England Biolabs). After staining, embryos were embedded in 1.5% agar/5% sucrose and frozen over dry ice. 20 μm transverse sections were cut using the Leica CM 1950 cryostat (Leica Microsystems), collected on microscope slides and mounted with 75% glycerol.

### Fluorescent In situ RNA Hybridization

Fluorescent in situ RNA hybridization was performed using the RNAScope Multiplex Fluorescent V2 Assay Kit (Advanced Cell Diagnostics; ACD) on 12 μm thick paraformaldehyde-fixed and agarose embedded cryosections according to manufacturer’s instructions with the following modification: slides were covered with parafilm for all 40°C incubations to maintain moisture and disperse reagents across the sections. The zebrafish *olig2*-C1, *nkx2.2a*-C2, *ptch2*-C2, *myrf*-C2, *cspg4*-C2, and *prdm8*-C3 transcript probes were designed and synthesized by the manufacturer and used at 1:50 dilutions. Transcripts were fluorescently labeled with Opal520 (1:1500), Opal570 (1:500) and Opal650 (1:1500) using the Opal 7 Kit (NEL797001KT; Perkin Elmer).

### Cold-active protease cell dissociation and FACs

24, 36, and 48 hpf *Tg(olig2:EGFP)* euthanized embryos were collected in 1.7 ml microcentrifuge tubes and deyolked in 100 μl of pre-chilled Ca free Ringers solution (116 mM NaCl, 2.6 mM KCl, 5 mM HEPES, pH 7.0) on ice. Embryos were pipetted intermittently with a p200 micropipettor for 15 minutes and left for 5 min. 500 μl of protease solution (10 mg/ml BI protease, 125 U/ml DNase, 2.5 mM EDTA, 1X PBS) was added to microcentrifuge tubes on ice for 15 min and embryos were homogenized every 3 min with a p100 micropipettor for 15 min. 200 μl of STOP solution (30% FBS, 0.8 mM CaCl_2_, 1X PBS) was then mixed into the tubes. Samples were then spun down at 400g for 5 min at 4°C and supernatant was removed. On ice, 1 ml of chilled suspension media (1% FBS, 0.8 mM CaCl_2_, 50 U/ml Penicillin, 0.05 mg/ml Streptomycin) was added to samples and then spun down again at 400g for 5 min at 4°C. Supernatant was removed, and 400 μl of chilled suspension media was added and solution was filtered through a 35 μm strainer into a collection tube. Cells were FAC sorted to distinguish EGFP^+^ cells using a MoFlo XDP100 cell sorter at the CU-SOM Cancer Center Flow Cytometry Shared Resource and collected in 1.7 ml FBS coated microcentrifuge tubes in 200 μl of 1X PBS.

### scRNA Sequencing

The Chromium Box from 10X Genomics was used to capture cells using Chromium Single Cell 3’ Reagent Kit part no. PN-1000075. Libraries were sequenced on the Illumina NovaSEQ6000 Instrument. FASTQ files were analyzed using Cell Ranger Software. 2174 (24h), 2555 (36h) and 3177 (48h) cells were obtained yielding a mean of 118,014 (24h), 65,182 (36h) and 96,053 (48h) reads per cell with a median of 1929 (24h), 1229 (36h) and 1699 genes identified per cell.

Raw sequencing reads were demultiplexed, mapped to the zebrafish reference genome (build GRCz11/danRer11) and summarized into gene-expression matrices using CellRanger (version 3.0.1). The resulting count matrices were further filtered in Seurat 3.1.0 (https://satijalab.org/seurat/) to remove cell barcodes with fewer than 250 detectable genes, more than 5% of UMIs derived from mitochondrial genes, or more than 50,000 UMIs (to exclude putative doublets). This filtering resulted in 6,489 single cells across all samples (1,952 from 24 hpf; 2,147 from 36 hpf; 2,390 from 48 hpf). After standard Seurat normalization, PCA was run using the 1,291 most variable genes. Next, dimensionality reduction was performed using Uniform Manifold Approximation and Projection (UMAP) on the first 15 principal components. Differential expression and marker gene identification was performed using MAST (https://doi.org/10.1186/s13059-015-0844-5).

### Immunohistochemistry

Larvae were fixed using 4% paraformaldehyde/1X PBS overnight at 4°C. Embryos were washed with 1X PBS, rocking at room temperature and embedded in 1.5% agar/5% sucrose, frozen over dry ice and sectioned in 20 or 15 µm transverse increments using a cryostat microtome. Slides were place in Sequenza racks (Thermo Scientific), washed 3×5 min in 0.1%Triton-X 100/1X PBS (PBSTx), blocked 1 hour in 2% goat serum/2% bovine serum albumin/PBSTx and then placed in primary antibody (in block) overnight at 4°C. The primary antibodies used include: rabbit anti-Sox10 (1:500; H.C. Park et al., 2005), mouse anti-Islet (1:500; DSH AB2314683), rat anti-BrdU (1:100; Abcam AB6326) or mouse JL-8 Living Colors (1:500; Clonetech 632380) to restore *Tg(olig2:EGFP)* fluorescence after RNA ISH. Sections were washed for 1 hours at room temperature with PBSTx and then incubated for 2 hr at room temperature with secondary antibodies at a 1:200 dilution in block. The secondary antibodies used include: AlexaFluor 488 anti-rabbit (Invitrogen A11008), AlexaFluor 588 anti-rabbit (Invitrogen A11011), AlexaFluor 647 anti-rabbit (Jackson Immuno. 111606144), AlexaFluor 488 anti-mouse (Life Tech., A11001), AlexaFluor 568 anti-mouse (Invitrogen A11004) and AlexaFluor 568 anti-rat (Invitrogen A11077). Sections were washed for 1 hr with PBSTx and mounted in VectaShield (Vector Laboratories).

### Imaging

Fixed sections of embryos and larvae were imaged on a Zeiss CellObserver SD 25 spinning disk confocal system (Carl Zeiss) or a Zeiss Axiovert 200 microscope equipped with a PerkinElmer spinning disk confocal system (Perkin Elmer Improvision). IHC cell counts were collected using a 20x objective (n.a. 0.8) and representative images were collected using a 40x oil immersion objective (n.a. 1.3). Wild-type 1 dpf larvae were positioned on top of a 2% agarose plate and imaged using a Leica M165FC dissection scope (Leica Microsystems) with a SPOT RT3 camera (SPOT Imaging). RNA ISH sections were imaged using differential interference contrast optics and a Zeiss AxioObserver compound microscope (Carl Zeiss). Cell counts and representative images were acquired at 40X (n.a. 0.75). Images are reported as extended z-projections or a single plan (RNA ISH) collected using Volocity (Perkin Elmer) or Zen (Carl Zeiss) imaging software. Image brightness and contrast were adjusted in Photoshop (Adobe) or ImageJ (National Institutes of Health).

## Data Quantification and Statistical Analysis

### Immunohistochemistry and RNA ISH

Quantifications of fluorescent cell numbers in transverse sections were performed by collecting confocal z-stacks of the entire section. Quantifications of RNA ISH cell numbers in transverse sections were performed by viewing the entire z-plane. Data for each embryo was collected from 10 consecutive trunk spinal cord sections and n represents the average number of cells per section in one embryo. All cell counts on sections were performed on blinded slides except Fig. 4A-D and Fig. 6E-F.

### Fluorescent RNA ISH

Quantification of fluorescent RNA ISH hybridization was carried out on z-projections collected at identical exposures. All quantification was performed in ImageJ Fiji using a custom script created by Karlie Fedder (available upon request). First, ten 0.5 μm z intervals were maximum z-projected and background was subtracted using a 2-rolling ball. The image was then thresholded by taking 2 standard deviations above the mean fluorescence intensity. A region of interest was drawn around the pMN domain or spinal cord and puncta were analyzed using the “Analyze Particles” feature with a size of 0.01-Infinity and circularity of 0.00-1.00. All thresholded puncta were inspected to ensure single molecules were selected. Puncta with an area of only 1 pixel were removed from the dataset. Data for each embryo was collected from 5 consecutives trunk spinal cord sections and n represents the average number of puncta in a region of interest per section in a single embryo.

### Statistical Analysis

We plotted all data and performed all statistical analyses in GraphPad Prism. All data are expressed as mean ± SEM. Normality was assessed with a D’Agostino and Pearson omnibus test. For statistical analysis, we used Student’s two-tailed t-test for all data with normal distributions or Mann-Whitney tests for non-normal data. Unless otherwise stated, all graphs represent data collected from one laboratory replicate, sampling fish from multiple crosses with no inclusion or exclusion criteria.

## Acknowledgements

We thank Christina Kearns for isolating cells for scRNA-seq and members of the Appel lab and the Section of Developmental Biology for discussions and advice. Cell sorting was performed by the University of Colorado Cancer Center Flow Cytometry Shared Resource, supported by the Cancer Center Support Grant (P30CA046934). scRNA-seq was performed by the University of Colorado Anschutz Medical Campus Genomics Shared Resource Core Facility, supported by the Cancer Center Support Grant (P30CA046934). Single cell RNA-sequencing and bioinformatics analysis was supported by a pilot award from the University of Colorado RNA Bioscience Initiative. The University of Colorado Anschutz Medical Campus Zebrafish Core Facility was supported by National Institutes of Health grant P30 NS048154.

## Data availability

All data are available within the manuscript.

